# Distinct Disruptions in CA1 and CA3 Place Cell Function in Alzheimer’s Disease Mice

**DOI:** 10.1101/2024.09.23.614631

**Authors:** Sanggeon Park, Mijeong Park, Eun Joo Kim, Jeansok J. Kim, Yeowool Huh, Jeiwon Cho

## Abstract

The hippocampus, a critical brain structure for spatial learning and memory, is susceptible to neurodegenerative disorders such as Alzheimer’s disease (AD). The APPswe/PSEN1dE9 (APP/PS1) transgenic mouse model is widely used to study the pathology of AD. Although previous research has established AD-associated impairments in hippocampal-dependent learning and memory, the neurophysiological mechanisms underlying these cognitive dysfunctions remain less understood. To address this gap, we investigated the activities of place cells in both CA1 and CA3 hippocampal subregions, which have distinct yet complementary computational roles. Behaviorally, APP/PS1 mice demonstrated impaired spatial recognition memory compared to wild-type (WT) mice in the object location test. Physiologically, place cells in APP/PS1 mice showed deterioration in spatial representation compared to WT. Specifically, CA1 place cells exhibited significant reductions in coherence and spatial information, while CA3 place cells displayed a significant reduction in place field size. Both CA1 and CA3 place cells in APP/PS1 mice also showed significant disruptions in their ability to stably encode the same environment. Furthermore, the burst firing properties of these cells were altered to forms correlated with reduced cognition. Additionally, the theta rhythm was significantly attenuated in CA1 place cells of APP/PS1 mice compared to WT. Our results suggest that distinct alteration in the physiological properties of CA1 and CA3 place cells, coupled with disrupted hippocampal theta rhythm in CA1, may collectively contribute to impaired hippocampal-dependent spatial learning and memory in AD.

## Introduction

Dementia affects more than 55 million people worldwide (1), with Alzheimer’s disease (AD) being the most prevalent form, accounting for 60-70% of all cases. The dorsal hippocampus (dHPC), a structure long been implicated in spatial navigation and memory (2, 3), is particularly affected in the early stage of AD. Most AD patients have difficulty forming hippocampal-dependent episodic and spatial memories (4, 5), attributed to the accumulation of amyloid beta (Aβ) plaques in the dHPC (6–8). To model the amyloid pathology and behavioral symptoms in animals, the amyloid precursor protein/presenilin 1 (APP/PS1) transgenic mice have been widely used in AD research (9–11). Previous investigations have established that APP/PS1 mice exhibit age-dependent hippocampal memory impairments when subjected to various behavioral tasks. For instance, studies employing the radial arm water maze have demonstrated deficits in mice between 3 to 5 months of age (11), while the Morris water maze and Barnes maze have shown impairments in mice starting at 6 months of age (12–14). Additionally, contextual fear conditioning has revealed deficits in 6 month-old mice (15), and passive avoidance tests have demonstrated impairments in mice between 6 to 12 months of age (13, 16) when compared to age-matched wildtype (WT) mice.

AD-related impairment of long-term potentiation (LTP) has also been studied in the hippocampal CA1 and CA3 regions in APP/PS1 mice (17, 18). Since several studies have shown that hippocampal LTP is associated with spatial memory and spatial firing patterns of place cells in rodents, altered hippocampal synaptic plasticity and memory in APP/PS1 mice may be related to aberrant firing patterns in CA1 and CA3 place cells. Hippocampal place cells are known to serve as a neural substrate for spatial learning and memory by encoding spatial locations in a given environment (2, 19–21). These cells display complex spike burst firing, which is implicated in synaptic plasticity and stability of place fields (22–24). In a familiar environment, most place cells have stable place fields for a long period of time (20, 25), while in the different environments, they changing firing rate (26) and/or firing location (27–29). These spatial firing patterns of place cells are postulated to provide a neurophysiological mechanism for encoding spatial location and memory retrieval.

Of the hippocampal subregions (CA1, CA2, CA3, CA4, and DG), prominent neuronal loss was found in the CA1 and CA3 of AD patients (30), suggesting the importance of the two regions in AD pathology. Since the CA1 and CA3 place cells are demonstrated to have distinct functions (31) (CA3 place cell showing stronger pattern completion tendency than CA1 place cells in similar environments), investigating changes in CA1 and CA3 place cells due to amyloid plaque deposition will confer deeper insight into understanding the hippocampal subregion specific pathophysiology of AD. Several studies report abnormal spatial coding of CA1 place cells in several AD rodent models (32–35) and one study reports pattern completion deficit of CA3 place cells in a mouse AD model (36). However, respective alterations in CA1 and CA3 place cell activities that lead to place field instability and spatial recognition memory deficit in AD remain unclear. To address this, we examined object location memory and spatial firing patterns of CA1 and CA3 place cells in 10-12 month-old APP/PS1 mice. Our study reports impairment in spatial learning and memory in the object location task and distinct yet significant alterations in activities of both the CA1 and CA3 place cells in APP/PS1 mice.

## Methods and Materials

### Animals

We used APP/PS1 transgenic mice from Jackson Laboratory, USA, (stock number 004462) with double mutations in the Swedish amyloid precursor protein (APPswe) and exon 9 (ΔE9) of the PSEN1 (presenilin 1; PS1) gene. Hemizygous mice were bred with B6C3F1/J mice. For the object location task, the experiment was carried out with both female and male APP/PS1 mice aged 10-12 months (N=9) and WT littermates (N=9) of the same age group. We used data obtained from 8 APP/PS1 and 9 WT mice after excluding an outlier. Exclusion was based on movement durations during the test phase: < mean of all movement durations (s) ± 2 standard deviations. Place cell recordings were performed on 10-12 month-old male APP/PS1 (N=13) and WT (N=13) mice. At least one place cell recording data that meets the inclusion criterion was obtained from the subjects and no subjects were excluded. All mice had free access to food and water and were maintained on a 12-hour light:dark cycle (lights on at 8 am). Mice were handled daily for 7 days (5 min/day) prior to experiments. All experimental procedures were approved and conducted in compliance with the Institutional Animal Care and Use Committee (IACUC) of Korea Institute of Science and Technology (Protocol Number: 2015-019). The sequence of subjects was randomized for all experiments and all experiments were carried out blinded to groups.

### Object location task

Both APP/PS1 and WT mice were habituated in the empty recording chamber for 2 days (10 min/day). After habituation, the mice were placed in the recording chamber with two identical objects and allowed to explore for 10 minutes (training session). One hour after training, the mice were returned to the recording chamber, where one of the objects had been moved to a new location, and they were allowed to explore for 5 minutes (memory test session). We recorded the time each mouse spent exploring each object during both training and memory test sessions. The exploration duration was analyzed by at least two investigators who were blinded to the group assignments.

### Surgical procedures for place cell recording

WT and APP/PS1 mice were anesthetized with Zoletil (30 mg/kg, i.p.) and placed on a stereotaxic instrument (David Kopf Instruments, USA) for chronic implantation surgery. A microdrive mounted with four tetrodes was implanted at specific coordinates for the dorsal hippocampal CA1 region (from bregma: AP -1.8 mm; ML - 1.5 mm; DV -0.6 mm from the brain surface) and for the CA3 region (AP -1.8 mm; ML -1.8 mm; DV -1.7 mm). This was done after drilling a hole above the right hippocampus. The microdrive was then secured onto the skull with dental cement and screws. Tetrodes (12.5 μm in diameter) were constructed by twisting four strands of polyimide-insulated nichrome microwires (Kanthal Precision Technology, Sweden) and fusing the insulation with heat. Each microwire tip was gold-plated to reduce the impedance to 300-500 kΩ (at 1 kHz).

### Place cell recording

After a seven-day recovery period, place cell recordings were performed using a Neuralynx Cheetah data acquisition system (Neuralynx, USA). Unit signals were amplified (10,000X), filtered (600 Hz-6 kHz), and sampled at 30.3 kHz. The mouse’s head angle and location were tracked and analyzed through two light-emitting diodes (LEDs) attached to the headstage, captured at 30 Hz by a ceiling-mounted video camera. Place cells were recorded in a black cylindrical open-field chamber (diameter 30 cm, height 12.7 cm) surrounded by a black curtain, with two spatial cues in a dimly lit room. The mice were gradually food-restricted until they reached 85% of their normal body weight to prepare them for a pellet chasing task. This task involved randomly dropping small food pellets (20 mg) into the recording chamber using a food pellet dispenser (Med-Associates Inc., USA). Two recording sessions, session A (SA) and session A’ (SA’), each lasting 20 minutes, were conducted with a 1-hour inter-session interval. A rectangular white cardboard (26 cm wide), covering a 90° arc, was mounted as a local cue inside of the cylinder for both sessions. White noise (80dB) was generated to minimize external noise interference.

### Place cell Analyses

Once neuronal signals were obtained, single units were isolated using cluster sorting software (SpikeSort 3D, Neuralynx, USA) and confirmed by inter-spike intervals (> 1 ms). Place cells were separated from the isolated single-units by mean firing rate of >0.2 Hz and the presence of bursts (high frequency spikes occurring within <15 ms with progressively decreasing amplitudes) as previously described (24, 37). Only place cell data with >0.2 Hz in both SA and SA’ were included in the analysis. After excluding place cells that does not meet our criterion, the following number of place cells were used for analyses: 35 cells out of a total of 50 cells in WT CA1, 25 out of a total of 39 cells in WT CA3, 38 cells out of a total of 51 cells in APP/PS1 CA1, and 28 out of a total of 52 cells in APP/PS1 CA3.

Rate maps (place fields) were created, represented by firing rates of each pixel, which were calculated by dividing the total number of spikes by the total time spent in each pixel, using 1 x 1 cm pixels smoothed with a 3×3 kernel. Pixels not visited for < 1 s during the 20-minute recording session were excluded from analysis. Spatial coherence was quantified by calculating the correlation between the firing rate of each pixel and the average firing rate of its eight neighboring pixels, assessing the uniformity of spatial firing patterns (20). Spatial information is calculated from neuronal signals to estimate how accurately the firing of a neuron can predict an animal’s location (38). Place field size (cm^2^) was calculated as the summed area of all pixels that had a firing rate higher than the overall firing rate. Spatial selectivity was quantified by calculating the logarithmic ratio of in-field to out-field firing rates, reflecting the specificity with which a neuron is dedicated to place coding [Spatial selectivity = log(In-field firing rate / Out-field firing rate)] (39). All data were analyzed using customized R-programs (40).

To measure stability, the similarity of place fields between two sessions was assessed by performing a pixel-by-pixel correlation, which was then transformed into Fisher’s Z score for parametric comparisons. Stability was further determined by identifying the degree of rotation between place maps that maximized the pixel-by-pixel cross-correlation, achieved through successive 5° rotations (22).

To provide a rough estimate of remapping, place cells were classified based on the maximum similarity achieved through specific degrees of rotation; cells maintaining maximum similarity within ±10 degrees between Session A (SA) and Session A’ (SA’) were labeled as “stay”, while those requiring greater rotations were identified as “remap”.

### Burst spike analysis

A burst was defined as two or more consecutive spikes with progressively decreasing peak amplitudes, each occurring within an inter-spike interval (ISI) of 15 ms (20, 22, 24). To analyze the distributions of the 1st and 2nd intra-burst intervals (intraBI) separately, we generated Cumulative Distribution Functions (CDFs) for each interval for both WT and APP/PS1 groups. Additionally, we calculated joint probability densities to assess the relationship between the two intervals, using Python’s Matplotlib and Seaborn libraries (41).

### Power spectral density

To compare the frequency characteristics of neuronal firing in the hippocampal CA3 and CA1 regions of APP/PS1 mice and WT mice, we analyzed the power spectral density of unit spike trains. This was done using Chronux (42) with specified parameters: moving window = [0.5, 0.05], tapers = [5, 9], and frequency pass range = [0.1, 100]. The relative PSD area was calculated by averaging across specific frequency bands: theta (6-10 Hz), low-gamma (25-60 Hz), and high-gamma (60-100 Hz) frequency bands (43, 44).

### Histology

After completing the recordings, the mice were overdosed with 2% Avertin and a small current (10-30 µA, 10 s) was passed through one of the four tetrode wires to mark the recording site. The mice were then transcardially perfused with 10% formalin, their brains were extracted, and further fixed in 10% formalin at room temperature. Coronal sections with a thickness of 50 μm were cut through the entire hippocampus region using a cryostat microtome (Microm, Germany) at -23°C. The brain slices were mounted on slides and stained with Cresyl Violet following the general Nissl staining procedure. The recording sites were examined to confirm whether the tetrodes had passed through the CA1 or CA3 pyramidal layer of the hippocampus.

### Thioflavin-S staining

Coronal brain sections, 50 μm thick, were washed three times in PBS solution. Subsequently, the sections were incubated in 1 mM Thioflavin-S, dissolved in 50% of ethanol, for 8 minutes. After incubation, the brain sections were rinsed twice with 80% ethanol and washed three more times with PBS solution. Images of the stained sections were captured using an Olympus BX50 microscope with a GFP filter at 40X magnification (13).

### Statistical Analysis

All statistical analyses were performed using SPSS statistics 27 (SPSS Inc., USA) and GraphPad Prism 8 (GraphPad Software Inc., USA). Group differences were evaluated using the Mann–Whitney U test. To analyze the distribution of “stay” and “remap” classifications of place cells between the two groups, the Chi-square test was employed. Additionally, the Kolmogorov–Smirnov test was used to analyze differences in cumulative rotational max similarity and the ISI density of IntraBI between groups.

## Results

### Distinct alterations in CA1 and CA3 place cell firing properties in APP/PS1 mice

#### Behavioral assessment in object location task

The study utilized the object location task to assess hippocampal-dependent spatial learning and memory in APP/PS1 mice. The experimental paradigm is depicted in Figure 1A. During the sample phase, both WT and APP/PS1 mice spent equal time exploring objects 1 and 2 (Figure 1B, sample phase). However, during the test phase, WT mice significantly favored exploring the relocated object 1 over the unmoved object 2 (Figure 1B, test phase), demonstrating intact spatial memory. Conversely, APP/PS1 mice showed no preference for the relocated object (Mann-Whitney U test; P=0.75; Figure 1B, test phase), indicating an impaired ability to recognize an object in a novel location. This observation aligns with previous findings (12–14) and confirms deficits in hippocampal-dependent spatial location memory in APP/PS1 mice.

**Figure 1.**
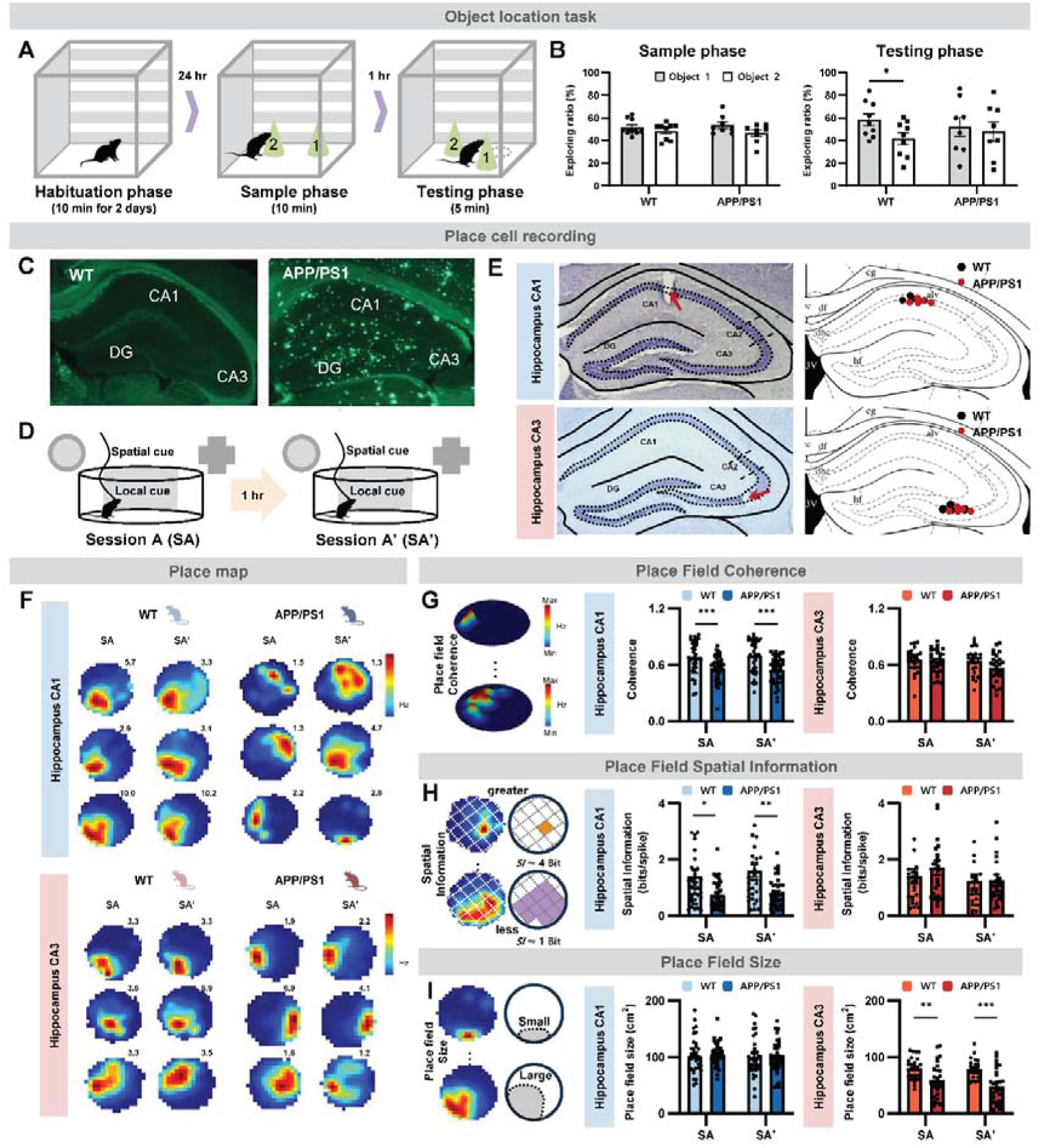
Impairments in object location memory and hippocampal place cell function in WT and APP/PS1 mice. (A) Schematic illustration of the object location task protocol. (B) Object preference during the sample and test phases for WT (N=9) and APP/PS1 (N=8) mice. Data are presented as mean±SEM. \**P* <0.05 (paired t-test). (C) Hippocampal amyloid plaque staining in WT and APP/PS1 mice using Thioflavin-S.. (D) Schematic of place cell recording experiments in hippocampal regions CA1 and CA3 for WT (N=10 mice, CA1=35 neurons, CA3=23 neurons) and APP/PS1 (N=9 mice, CA1=38 neurons, CA3=28 neurons). (E) Histological samples showing CA1 and CA3 place cell recording sites. Black dots indicate recordings from WT mice, while red dots indicate recordings from APP/PS1 mice. (F) Representative place maps of CA1 and CA3 from WT and APP/PS1 mice in SA and SA’ sessions.. (G) Place field coherence for CA1 and CA3 place cells from WT and APP/PS1 mice recorded during SA and SA’ sessions. (H) Spatial information content of place fields for CA1 and CA3 in WT and APP/PS1 mice derived from SA and SA’ sessions. (I) Size of place fields in CA1 and CA3 for WT and APP/PS1 mice, assessed during SA and SA’ sessions. (G-I) Data are presented as mean±SEM. Statistical significance assessed by Mann-Whitney U test; *P <0.05, **P <0.01, ***P <0.001..

#### Neurophysiological observations and place cell activity

To explore neurophysiological alterations, we recorded the activity of dorsal CA1 and CA3 pyramidal neurons in freely behaving WT and APP/PS1 mice aged 10-12 months (WT N=10, APP/PS1 N=9). Significant amyloid plaque deposition was observed throughout the dorsal hippocampal formation in APP/PS1 mice compared to age-matched WT mice (Figure 1C). Mice were exposed to the same environment with an hour interval between sessions (SA: session A, SA’: session A’) to assess the stability of CA1 and CA3 place cells in encoding and recalling similar environments (Figure 1D). Our analysis included 60 place cells from WT mice and 66 place cells from APP/PS1 mice, all exhibiting firing rates above 0.2 Hz (CA1=35, CA3=25 for WT; CA1=38, CA3=28 for APP/PS1). Place cell activities of the WT and APP/PS1 mice were obtained from similar locations (Figure 1E). Sample place maps from CA1 and CA3 place cells in sessions SA and SA’ are shown in Figure 1F.

#### Quantitative analysis of place cell properties

The analysis suggested that the integrity of APP/PS1 CA1 place cells was more compromised than that of CA3 place cells. Measures of coherence and spatial information for APP/PS1 CA1 place cells were significantly reduced in both sessions compared to WT (Mann-Whitney U test; coherence: SA P=0.001, SA’ P=0.001; spatial information: SA P=0.015, SA’ P=0.002; Figures 1G, H). The size of place fields, however, did not differ significantly from WT (Mann-Whitney U test; SA P=0.42, SA’ P=0.93; Figure 1I). In contrast, coherence and spatial information of APP/PS1 CA3 place cells were comparable to WT (Mann-Whitney U test; coherence: SA P=0.789, SA’ P=0.057; spatial information: SA P=0.206, SA’ P=0.569; Figures 1G, H), but their place field size was significantly reduced (Mann-Whitney U test; SA P=0.003, SA’ P<0.001; Figure 1I).

#### Consistency and variability in place cell properties

Although APP/PS1 CA1 place cells showed significant reductions in coherence and spatial information compared to WT, other parameters such as mean firing rate, maximum firing rate, firing rate within or outside a place field, and spatial selectivity remained unchanged in the initial SA session (Table 1). However, upon re-exposure to the same environment, APP/PS1 CA1 place cells significantly reduced maximum firing rate (Mann-Whitney U test; P=0.043), firing rate within a place field (Mann-Whitney U test; P=0.026), and spatial selectivity (Mann-Whitney U test; P=0.036) compared to the WT (Table 1). No significant differences were observed in these parameters between APP/PS1 and WT CA3 place cells (Table 1).

**Table 1.** Comparison of firing properties between WT and APP/PS1 place cells in CA1 and CA3. Mann-Whitney U test between WT and APP/PS1. *P<0.05.

| Parameters | Session | CA1 |  | CA3 |  |
| --- | --- | --- | --- | --- | --- |
|  |  | WT | APP/PS1 | WT | APP/PS1 |
| Spikes | SE1 | 1365.2 ±203.3 | 1035.5 ±121.8 | 1172.0 ±248.4 | 1326.9 ±190.3 |
|  | SE2 | 1677.2 ±233.1 | 1187.2 ±148.8 | 1332.8 ±405.9 | 1075.1 ±172.8 |
| Mean Firing rate (Hz) | SE1 | 1.13 ±0.16 | 0.86 ±0.10 | 0.97 ±0.20 | 1.10 ±0.15 |
|  | SE2 | 1.39 ±0.19 | 0.98 ±0.12 | 1.11 ±0.33 | 0.89 ±0.14 |
| Max firing rate (Hz) | SE1 | 4.88 ±0.67 | 2.90 ±0.21 | 3.37 ±0.42 | 3.54 ±0.64 |
|  | SE2 | 5.14 ±0.72 | 2.93 ±0.25 * | 3.31 ±0.81 | 2.64 ±0.48 |
| In-field firing rate (Hz) | SE1 | 3.08 ±0.44 | 1.88 ±0.15 | 2.21 ±0.34 | 2.25 ±0.33 |
|  | SE2 | 3.48 ±0.49 | 2.02 ±0.19 * | 2.34 ±0.62 | 1.87 ±0.27 |
| Out-field firing rate (Hz) | SE1 | 0.37 ±0.06 | 0.39 ±0.06 | 0.29 ±0.07 | 0.28 ±0.04 |
|  | SE2 | 0.50 ±0.09 | 0.46 ±0.08 | 0.45 ±0.17 | 0.20 ±0.04 |
| Selectivity | SE1 | 1.04 ±0.03 | 0.87 ±0.05 | 0.99 ±0.04 | 0.96 ±0.04 |
|  | SE2 | 0.99 ±0.03 | 0.82 ±0.05 | 0.94 ±0.04 | 0.99 ±0.04 |

Despite less pronounced differences between APP/PS1 and WT CA3 place cells compared to CA1, the ability of APP/PS1 CA3 place cells to stably encode the same environment was significantly compromised. Unlike WT CA3 place cells, which showed consistent properties across sessions, APP/PS1 CA3 place cells displayed significant differences in mean number of spikes (Paired t-test; P=0.048), mean firing rate (P=0.048), maximum firing rate (t-test; P=0.012), in-field firing rate P=0.017), out-field firing rate (P=0.018), coherence (P=0.003), spatial information P=0.031), and place field size (P=0.011) between sessions SA and SA’ (Table 2). Notably, the properties of APP/PS1 CA1 place cells remained consistent across sessions (Table 2), indicating a relative preservation of their capability to encode the same environment despite some properties being significantly diminished compared to WT CA1 place cells.

**Table 2.** Comparison of firing properties between sessions SA and SA’ for WT and APP/PS1 place cells in CA1 and CA3. Paired t-test between SA and SA’. *P<0.05.

| Parameters | Group | CA1 |  | CA3 |  |
| --- | --- | --- | --- | --- | --- |
|  |  | SE1 | SE2 | SE1 | SE2 |
| Spikes | WT | 1365.2 ±203.3 | 1677.2 ±233.1 | 1172.0 ±248.4 | 1332.8 ±405.9 |
|  | APP/PS1 | 1035.5 ±121.8 | 1187.2 ±148.8 | 1326.9 ±190.3 | 1075.1 ±172.8 * |
| Mean Firing rate (Hz) | WT | 1.13 ±0.16 | 1.39 ±0.19 | 0.97 ±0.20 | 1.10 ±0.15 |
|  | APP/PS1 | 0.86 ±0.10 | 0.98 ±0.12 | 1.11 ±0.33 | 0.89 ±0.14 * |
| Max firing rate (Hz) | WT | 4.88 ±0.67 | 5.14 ±0.72 | 3.37 ±0.42 | 3.54 ±0.64 |
|  | APP/PS1 | 2.90 ±0.21 | 2.93 ±0.25 | 3.31 ±0.81 | 2.64 ±0.48 * |
| In-field firing rate (Hz) | WT | 3.08 ±0.44 | 3.48 ±0.49 | 2.21 ±0.34 | 2.25 ±0.33 |
|  | APP/PS1 | 1.88 ±0.15 | 2.02 ±0.19 | 2.34 ±0.62 | 1.87 ±0.27 * |
| Out-field firing rate (Hz) | WT | 0.37 ±0.06 | 0.50 ±0.09 | 0.29 ±0.07 | 0.28 ±0.04 |
|  | APP/PS1 | 0.39 ±0.06 | 0.46 ±0.08 | 0.45 ±0.17 | 0.20 ±0.04 * |
| Selectivity | WT | 1.04 ±0.03 | 0.99 ±0.03 | 0.99 ±0.04 | 0.96 ±0.04 |
|  | APP/PS1 | 0.87 ±0.05 | 0.82 ±0.05 | 0.94 ±0.04 | 0.99 ±0.04 |

### Impaired stability of spatial representation in both CA1 and CA3 place cells of APP/PS1 mice

#### Analysis of place field stability

To evaluate the stability of place fields in CA1 and CA3 place cells between WT and APP/PS1 mice in a familiar environment, we analyzed several characteristics of the place fields from sessions SA and SA’. The methods used to assess place field stability are detailed in Figure 2A. We calculated the pixel-by-pixel cross-correlation between place fields from SA and SA’. Rotational similarity was assessed by computing cross-correlation values when the SA’ place field was rotated in 5° increments, identifying the maximum similarity as the highest cross correlation value achieved when the SA’ place field was rotated 5° clockwise. Additionally, we measured the distance between the peaks of place fields from SA and SA’.

**Figure 2.**
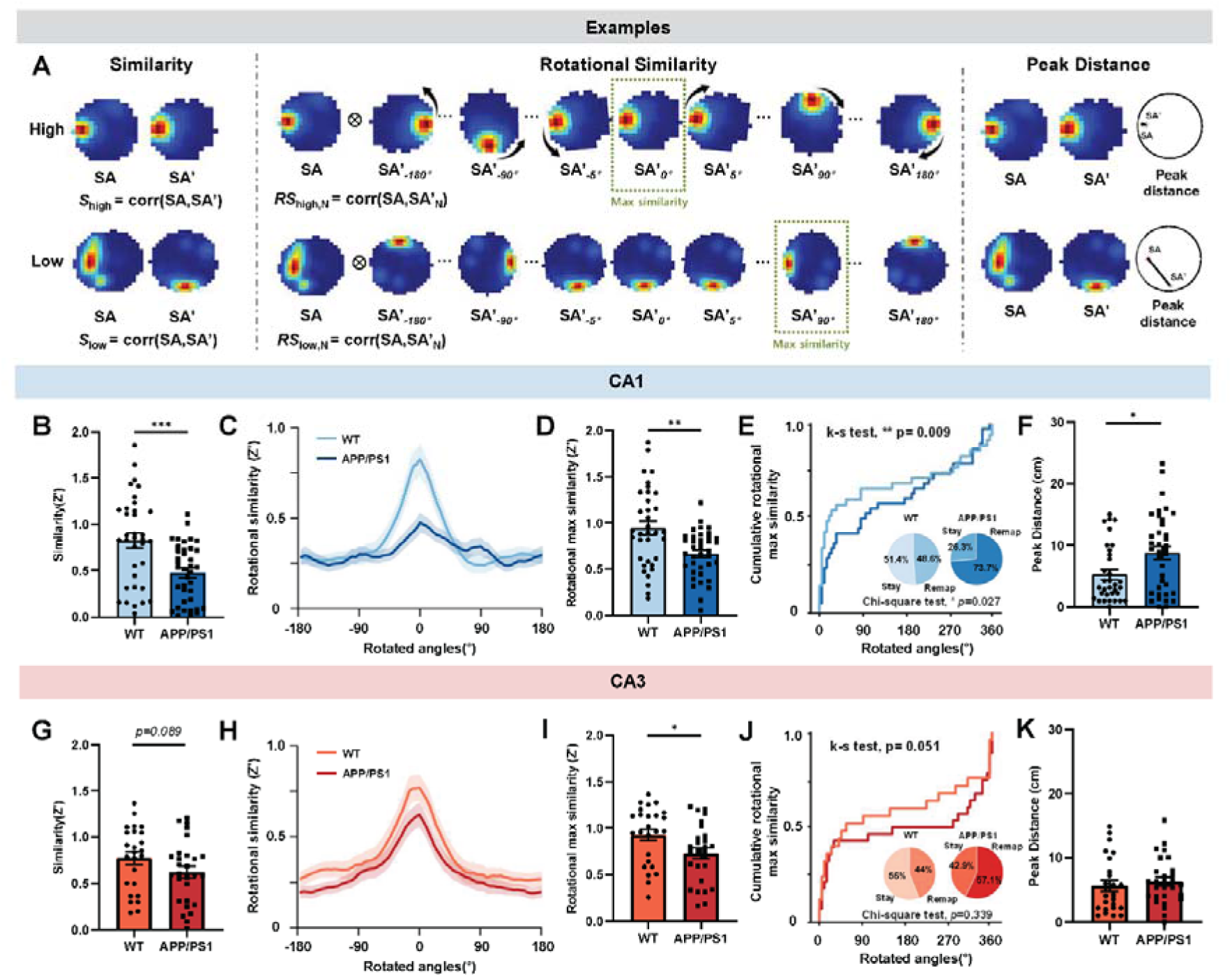
Reduced CA1 and CA3 place field similarities in APP/PS1 mice between environments SA and SA’. (A) Schematic drawing of place cell similarity tests used to assess the stability of place maps in environments SA and SA’: similarity, rotational similarity, and distance between peaks. (B) CA1 place field similarity (Z-transformed) between environments SA and SA’ for WT and APP/PS1 mice. (C) Rotational similarity (Z-transformation) between environments SA and SA’ for WT and APP/PS1 mice. (D) Maximum rotational similarity (Z-transformed) of CA1 place fields for WT and APP/PS1 mice. (E) Cumulative maximum rotational similarity of CA1 place fields for WT and APP/PS1 mice. Kolmogorov-Smirnov test, P=0.009. Relative ratio of place fields that remained stable (within ±10°) or remapped. (F) Distance between peak firing locations of CA1 place cells in SA and SA’ for WT and APP/PS1 mice. (G) CA3 place field similarity (Z-transformed) between environments SA and SA’ for WT and APP/PS1 mice. (H) Rotational similarity (Z-transformed) of CA3 place fields for WT and APP/PS1 mice. (I) Maximum rotational similarity (Z-transformed) of CA3 place fields for WT and APP/PS1 mice. (J) umulative maximum rotational similarity of CA3 place fields for WT and APP/PS1 mice. Kolmogorov-Smirnov test, P=0.051. Relative ratio of WT and APP/PS1 CA1 place fields that remained stable (within ±10°) or remapped. (K) Distance between peak firing locations of CA3 place cells in SA and SA’ for WT and APP/PS1 mice. (B, D, F, G, H, K) Bar graphs are expressed as mean±SEM. Statistical significance assessed using the Mann-Whitney U test; *P <0.05, **P <0.01, ***P <0.001.

#### Comparison of CA1 place field stability between WT and APP/PS1

Place field similarity between SA and SA’ was significantly lower in APP/PS1 CA1 place cells compared to WT (Mann-Whitney U test, p<0.001; Figure 2B). This trend extended to the rotational similarity test, where higher rotational similarity values indicated greater similarity, typically peaking around zero degrees (Figure 2C). The maximum similarity value was also significantly lower in APP/PS1 CA1 place cells than in WT (Mann-Whitney U test, P=0.0027; Figure 2D), as evidenced in the cumulative rotational similarity test (K-S test, P=0.009; Figure 2E). A significantly greater number of APP/PS1 CA1 place maps were categorized as remapped (rotation greater than 10°) compared to WT (Chi-square test, P=0.027; Figure 2E). The distance between the peaks of SA and SA’ was also significantly greater in APP/PS1 CA1 place maps compared to WT (Mann-Whitney U test, P=0.0271; Figure 2F), indicating a greater peak shift between sessions in APP/PS1 CA1 place maps.

#### Relative stability of APP/PS1 CA3 place cells

Compared to APP/PS1 CA1 place maps, the stability of APP/PS1 CA3 place maps was relatively preserved. Only the rotational maximum similarity of APP/PS1 CA3 place maps was significantly lower than that of WT CA3 (Mann-Whitney U test, P=0.0296; Figure 2I). Other parameters such as overall similarity, cumulative rotational similarity, the ratio of remapped place cells, and the distance between peaks of place maps between APP/PS1 and WT CA3 place cells showed no significant differences (Figure 2G, H, J, K).

### Abnormal bursting patterns in CA1 and CA3 of APP/PS1 mice

#### Role of hippocampal burst firing

Our previous studies have established that hippocampal burst firing plays a critical role in hippocampus-dependent spatial learning and memory as well as in the stability of place cells (22, 24). Given that APP/PS1 mice exhibit deficits in object location memory and place cell stability (Figures 1 and 2), we examined whether burst firing patterns in CA1 and CA3 were similarly altered in APP/PS1 mice. A burst was defined as two or more consecutive spikes with progressively decreasing amplitudes within an inter-spike interval (ISI) of 15 ms, as previously described (20, 22, 24). The properties of bursts analyzed are depicted in Figure 3A.

**Figure 3.**
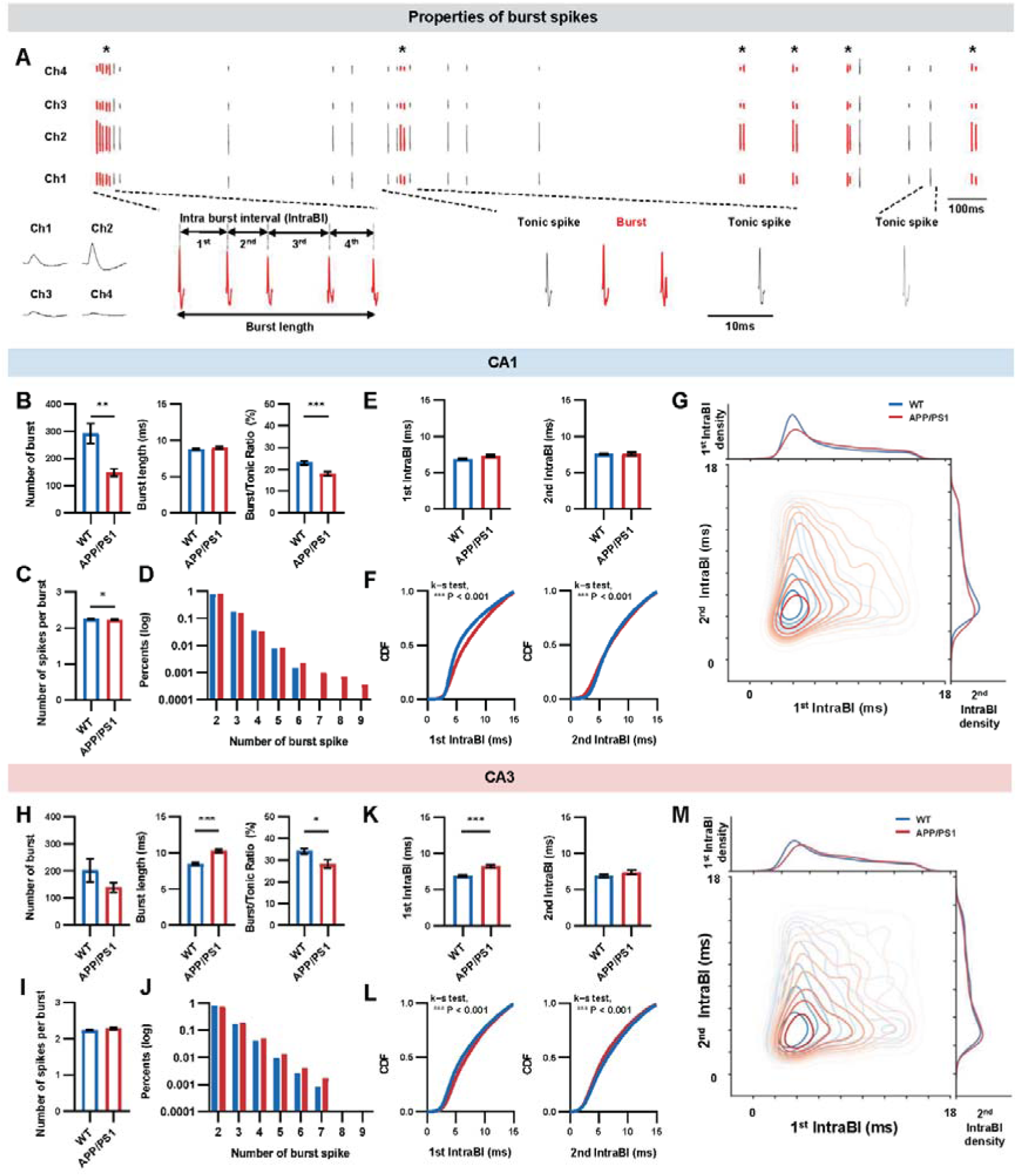
Altered burst firing properties of APP/PS1 CA1 and CA3 place cells compared to the WT. (A) Sample spike trains obtained from a tetrode, highlighting analyzed burst properties. Spikes colored in red indicate burst spikes, with asterisks marking the occurrence of bursts. (B) Number of bursts, burst length, and ratio of burst to tonic spike in WT and APP/PS1 CA1 place cells. (C) Number of bursts, burst length, and ratio of burst to tonic spikes in WT and APP/PS1 CA1 place cells. (D) Percentage of bursts composed of different burst spike counts in WT and APP/PS1 CA1 place cells. (E) Length of the first and second intra-burst-interval (IntraBI) in WT and APP/PS2 CA1 place cells. (F) Cumulative distribution function of the first and second IntraBI lengths for WT and APP/PS2 CA1 place cells. (G) Joint probability distribution of the first and second IntraBI in WT and APP/PS1 CA1 place cells. (H) Number of bursts, burst length, and ratio of burst to tonic spikes in WT and APP/PS1 CA3 place cells. (I) Number of burst spikes per burst in WT and APP/PS1 CA3 place cells. (J) Percentage of bursts composed of different burst spike numbers in WT and APP/PS1 CA3 place cells. (K) Length of the first and second IntraBI in WT and APP/PS2 CA3 place cells. (L) Cumulative distribution function of the first and second IntraBI lengths for WT and APP/PS2 CA3 place cells. (M) Joint probability distribution of the first and second IntraBI in WT and APP/PS1 CA3 place cells. (B-E, H-K) Bar graphs are presented as mean±SEM. Statistical significance assessed using the Mann-Whitney U test; *P <0.05, **P <0.01, ***P <0.001.

#### Distinct burst firing patterns

The burst firing patterns of APP/PS1 CA1 and CA3 place cells were altered in distinct ways compared to WT mice. APP/PS1 CA1 place cells showed a significantly reduced number of burst occurrences (Mann Whitney U test, P=0.008) and a lower burst/tonic ratio (P<0.001) compared to WT (Figure 3B). Additionally, the mean number of bursts spikes composing a burst was significantly reduced in APP/PS1 CA1 place cells compared to WT (APP/PS1: 2.23±0.01, WT: 2.25±0.01; Mann Whitney U test, P=0.012; Figure 3C), though some bursts in APP/PS1 CA1 place cells had more spikes than those in WT (Figure 3D). In contrast, APP/PS1 CA3 place cells did not show significant differences in the number of burst occurrences (Mann Whitney U test, P=0.656; Figure 3H) or the number of spikes per burst (P=0.203; Figure 3I) compared to WT. However, APP/PS1 CA3 place cells exhibited significantly longer burst lengths (P<0.001) and a lower burst/tonic ratio (P=0.021) relative to WT (Figure 3H).

#### Intervals between burst spikes (IntraBI)

The intervals between burst spikes in CA1 and CA3 place cells also showed variable patterns; the mean length of the first and second IntraBI in APP/PS1 CA1 place cells were similar to those in WT (Mann Whitney U test, 1^st^ IntraBI P=0.0718, 2^nd^ IntraBI P=0.7597; Figure 3E). However, the first IntraBI in APP/PS1 CA3 place cells was significantly longer than in WT (P<0.001; Figure 3K). Although the mean lengths of IntraBIs were generally preserved, the cumulative distribution of IntraBIs in APP/PS1 CA1 and CA3 place cells differed significantly from WT (K-S test, P<0.001 for both CA1 and CA3 1^st^ and 2^nd^ IntraBI; Figures 3F and L), indicating that collectively, the lengths of APP/PS1 IntraBIs are altered compared to WT. The joint probability analyses of the first and second IntraBIs qualitatively showed that the contour lines of both APP/PS1 CA1 and CA3 place cells are more dispersed than those of WT, indicating a deterioration in the integrity of the burst spikes composing bursts (Figures 3G and M).

#### Implications of altered burst firing

These findings align with previous reports of significant alterations in various burst properties in mouse models with memory deficits (22, 24). The altered quality and quantity of burst firing in APP/PS1 CA1 and CA3 place cells, which diverge in specific patterns from WT, suggesting that these abnormalities may contribute to the reduced recognition of relocated objects in APP/PS1 mice, as burst firing is known to have greater information content than tonic firing (23). This underscores the potential role of abnormal bursting patterns in the cognitive deficits observed in AD models.

### Reduced theta power of APP/PS1 CA1 place cells

Hippocampal theta, low, and high gamma rhythms have been implicated in spatial learning and memory (43, 45, 46). To explore potential changes in hippocampal rhythms underlying spatial memory deficits, we assessed the power spectral density of theta (6-10 Hz), low gamma (25-60 Hz), and high gamma (60-100 Hz) in CA1 and CA3 place cells of APP/PS1 and WT groups (43, 44). Our results were consistent with previous studies (32, 47, 48), showing that the theta power in APP/PS1 CA1 place cells was significantly lower than in WT (Mann Whitney U test, P=0.020; Figure 4A). However, the gamma rhythms in these cells did not differ significantly from those in WT (low gamma P=0.895, high gamma P=0.125; Figure 4A). Conversely, none of the rhythms in APP/PS1 CA3 place cells showed significant differences from WT (theta P=0.498, low gamma P=0.346, high gamma P=0.208; Figure 4B). These results indicate that while the theta rhythm is significantly attenuated in the CA1 of APP/PS1 mice, both low and high gamma rhythms remain preserved. In contrast, hippocampal rhythms in the CA3 of APP/PS1 mice are relatively intact. This suggests that the function of the APP/PS1 CA1 may be more adversely affected than that of APP/PS1 CA3, contributing to the spatial memory deficits observed in these mice. These findings highlight the differential impact of Alzheimer’s pathology on specific hippocampal subregions and their associated rhythmic activities.

**Figure 4.**
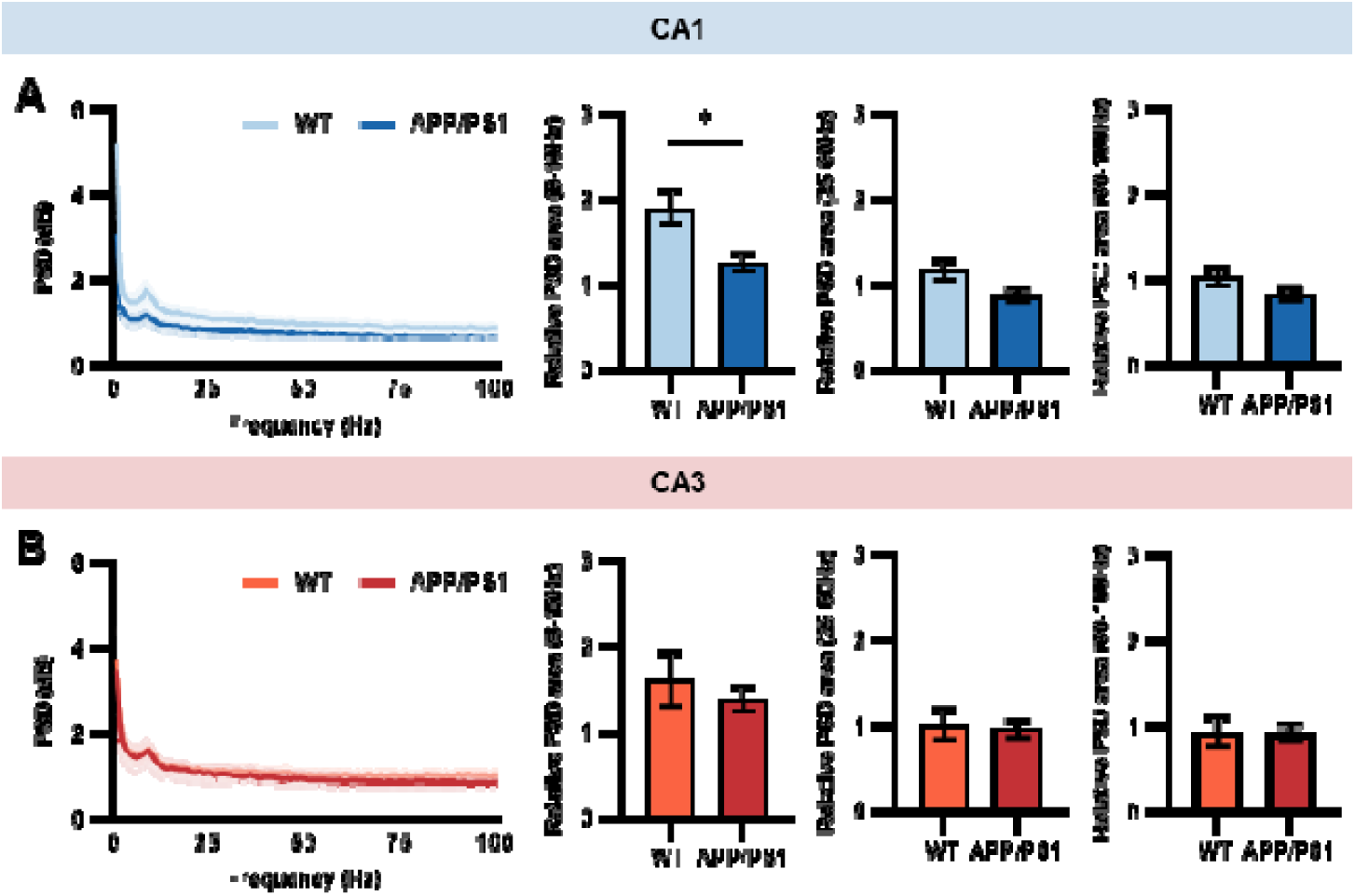
Hippocampal rhythm analysis of WT and APP/PS1 CA1 and CA3 place cells. (A) Power spectral density (PSD) analysis of CA1 place cells in WT and APP/PS1 mice, including comparisions of the PSD area across the theta (6-10 Hz), low gamma (25-60 Hz), and high gamma (60-100 Hz) frequency ranges. (B) PSD analysis of CA3 place cells in WT and APP/PS1 mice, similarly comparing the PSD area across the theta, low gamma, and high gamma frequency ranges. (A, B) Bar graphs represent data as mean±SEM. Statistical significance assessed using the Mann-Whitney U test; *P <0.05.

## Discussion

The present study investigates the neurophysiological mechanisms underlying hippocampal-dependent spatial learning and memory deficits in AD using the APP/PS1 mouse model. Our findings indicate that APP/PS1 mice exhibit deficits in object location memory, which correspond with alterations in CA1 and CA3 place cell firing properties. Specifically, APP/PS1 CA1 place cells demonstrated reduced coherence and spatial information, while APP/PS1 CA3 place cells exhibited significantly smaller place fields. Both CA1 and CA3 place cells in APP/PS1 mice showed a compromised ability to stably represent the same environment, suggesting that altered firing properties and less stable spatial representations may contributed to difficulties in recognizing a familiar environment.

Our findings are consistent with numerous previous studies reporting impaired hippocampal-dependent learning and memory in APP/PS1 mice (11, 12, 15, 16, 22, 49). We also confirmed declines in spatial recognition memory in APP/PS1 mice, as evidenced by their lack of preference for an object moved to a new location during the memory test session. This behavioral deficit is likely due to impaired hippocampal functions necessary for recognizing the previous locations of objects within the same environment (50–52).

While both CA1 and CA3 place cells in APP/PS1 mice displayed altered firing properties, the patterns of these alterations differed. CA1 place cells showed significantly lower coherence and spatial information (Figure 1G, H), suggesting a substantial reduction in their spatial encoding capabilities compared to WT. Conversely, CA3 place cells maintained their coherence and spatial information but texhibited significantly smaller place fields compared to WT. Given that the size of a place field is generally proportional to the extent of space it encodes (53), the reduced place field size in APP/PS1 CA3 place cells may inadequately represent the area of the recording chamber.

The ability of APP/PS1 CA1 and CA3 place cells to stably represent the same environment in successive sessions (SA and SA’) was significantly impaired compared to WT, as detailed in the place field stability analyses (Figure 2). The stability of place fields was more markedly diminished in APP/PS1 CA1 than in APP/PS1 CA3, indicating a more severe functional impairment in CA1 place cells. This finding aligns with clinical observations noting that while AD patients exhibit significant neuronal loss in both the CA1 and CA3 regions, the most pronounced decrease occurs in the CA1 area (30). Further supporting this, another study reported that CA1, compared to CA3 or DG, showed the highest number of neurofibrillary tangles (NFTs) and amyloid plaques (54). It has also been documented that NFTs initially accumulate in the CA1 region and subsequently affect the subiculum, CA2, CA3, and DG as AD progresses (55). Consequently, the function of APP/PS1 CA3 place cells may be relatively spared in the early stages of AD progression.

Burst firing, known for its high information content (23) and potential to multiplex signals (56), plays crucial role in spatial memory. Animals with place memory deficits often exhibit significant alterations in burst spiking patterns in CA1 place cells, such as a reduced number of bursts and elongated intervals between burst spikes (IntraBI) (22, 24). Our study found similar alterations in burst firing properties in APP/PS1 mice. Specifically, APP/PS1 CA1 place cells displayed a significantly reduced number of bursts, whereas APP/PS1 CA3 place cells had a significantly extended first IntraBI compared to WT. Additionally, both APP/PS1 CA1 and CA3 place cells showed a notable reduction in the ratio of bursts, and their cumulative distribution functions for the first and second IntraBIs were significantly distinct from those of WT cells. These changes suggest that altered burst firing properties in these place cells may underlie the observed impairments in place memory.

While it is not yet clear what molecular and circuit level changes are causing burst firing property alteration in the APP/PS1 CA1 and CA3 place cells, hippocampal burst probability can be modulated by various factors such as extracellular Ca^2+^ concentration (57), cell-type specific afferent activity (19, 58), Ca^2+^-dependent channels (e.g. L-, N-, T-, P/Q type), AMPA (α-amino-3-hydroxy-5-methyl-4-isoxazolepropionic acid) receptors, and NMDA (N-methyl-D-aspartate) receptors (59–61). There is evidence that Aβ could induce dysregulation of Ca^2+^ signaling in the hippocampus by affecting Ca^2+^-dependent channels and glutamate receptors (AMPA and NMDA) (62–64). For example, Aβ increases Ca^2+^ entry through voltage-gated Ca^2+^ channels (N-, T-, L-type) and NMDA receptors (62, 65–68) while reducing presynaptic L-type and P/Q-type Ca^2+^ channels activity and AMPA current (64, 69–71), resulting in neurotoxicity and reduced synaptic activity. Additionally, we have shown that levels of CaMKII proteins, key molecules for LTP induction in the hippocampus (72, 73), strongly correlate with extended lengths of inter-burst intervals and IntraBIs in the hippocampus (22, 24). Earlier studies have reported Aβ-induced LTP impairment in CA1 and overall reductions in phospho-CaMKII protein levels in both the CA1 and CA3 regions of the hippocampus in APP/PS1 mice (74). These physiological and molecular changes may collectively contribute to altering burst spiking patterns in both the CA1 and CA3 place cells of APP/PS1 mice.

Our study’s analysis of hippocampal rhythms demonstrated that most rhythms are preserved despite significant Aβ accumulation in the age range tested (10-12 months). Notably, only the theta rhythm in the CA1 was significantly reduced in APP/PS1 mice compared to the WT, while both low and high gamma rhythms remained intact. This result aligns with findings from studies in AD animal models that report a reduction in theta power in the hippocampus (32, 47, 48). Additionally, a study involving the 5xFAD mouse model—an Aβ overexpression model of AD— reported that gamma rhythm deficits are only evident in older mice (>12 months) and remain unaltered from 6 to 12 months despite significant Aβ accumulation (75). This suggests that gamma rhythms may be preserved until later stages of AD and are not likely the cause of spatial memory deficits observed in earlier stages. In the CA3 region of APP/PS1 mice, all rhythms were intact and showed no differences from those in WT, supporting the notion that the functions of CA1 may be more adversely affected by Aβ accumulation than those of CA3.

The relatively preserved physiological activities of the APP/PS1 CA3 place cells partly explain different dysfunctions that develop with the progression of AD. Since the CA3 has extensive recurrent collaterals, an auto-associative network, it has been implicated in one-step learning (76). The CA3 place cells was shown to have greater pattern completion, generalization, tendency than the CA1 place cells (31). Patients with mild cognitive impairment (MCI) displayed decrease in pattern separation rate, but similar pattern completion rates with healthy adults, while AD patients showed disruption in both pattern completion and pattern separation (77). This suggests that the intact pattern completion ability in MCI patients may be due to relatively preserved CA3 function.

Overall, our study provides new evidence for understanding the hippocampal neuronal pathophysiology and circuit mechanisms involved in spatial memory impairments in AD by demonstrating abnormal APP/PS1 place cell activities in both CA1 and CA3 regions. The different CA1 and CA3 place cells’ alteration patterns in firing activities, place field stability, burst patterns, and hippocampal rhythm may collectively be responsible for the impaired spatial stability and memory deficits observed in APP/PS1 mice.

## Acknowledgments

This work was supported by the National Research Foundation of Korea (NRF) grants funded by the Ministry of Science and ICT [NRF-2022M3E5E8018421 (J.C.) and NRF-2022R1A2C2009265 (J.C.)] and by NIH grants AG067008 (EJK) and MH099073 (JJK). The authors declare no competing financial interests.

## References

1. WHO, Dementia Fact sheet. World Health Organization (2022).

2. E. I. Moser, E. Kropff, M. B. Moser, Place cells, grid cells, and the brain’s spatial representation system. Annu Rev Neurosci 31, 69–89 (2008).

3. J. O’keefe, L. Nadel, The hippocampus as a cognitive map. Oxford University Press (1978).

4. E. Arnaiz, O. Almkvist, Neuropsychological features of mild cognitive impairment and preclinical Alzheimer’s disease. Acta Neurol Scand Suppl 179, 34–41 (2003).

5. R. Sperling, Functional MRI studies of associative encoding in normal aging, mild cognitive impairment, and Alzheimer’s disease. Ann N Y Acad Sci 1097, 146–155 (2007).

6. F. Cacucci, M. Yi, T. J. Wills, P. Chapman, J. O’Keefe, Place cell firing correlates with memory deficits and amyloid plaque burden in Tg2576 Alzheimer mouse model. Proceedings of the National Academy of Sciences of the United States of America 105, 7863–7868 (2008).

7. P. F. Chapman et al., Impaired synaptic plasticity and learning in aged amyloid precursor protein transgenic mice. Nat Neurosci 2, 271–276 (1999).

8. D. J. Selkoe, Alzheimer’s disease is a synaptic failure. Science 298, 789–791 (2002).

9. D. R. Borchelt et al., Accelerated amyloid deposition in the brains of transgenic mice coexpressing mutant presenilin 1 and amyloid precursor proteins. Neuron 19, 939–945 (1997).

10. R. S. Reiserer, F. E. Harrison, D. C. Syverud, M. P. McDonald, Impaired spatial learning in the APPSwe + PSEN1DeltaE9 bigenic mouse model of Alzheimer’s disease. Genes, brain, and behavior 6, 54–65 (2007).

11. A. Volianskis, R. Kostner, M. Molgaard, S. Hass, M. S. Jensen, Episodic memory deficits are not related to altered glutamatergic synaptic transmission and plasticity in the CA1 hippocampus of the APPswe/PS1deltaE9-deleted transgenic mice model of ss-amyloidosis. Neurobiology of aging 31, 1173–1187 (2010).

12. D. A. Gimbel et al., Memory impairment in transgenic Alzheimer mice requires cellular prion protein. The Journal of neuroscience : the official journal of the Society for Neuroscience 30, 6367–6374 (2010).

13. S. Jo et al., GABA from reactive astrocytes impairs memory in mouse models of Alzheimer’s disease. Nature medicine 20, 886–896 (2014).

14. R. Lalonde, H. D. Kim, J. A. Maxwell, K. Fukuchi, Exploratory activity and spatial learning in 12-month-old APP(695)SWE/co+PS1/DeltaE9 mice with amyloid plaques. Neuroscience letters 390, 87–92 (2005).

15. M. Kilgore et al., Inhibitors of class 1 histone deacetylases reverse contextual memory deficits in a mouse model of Alzheimer’s disease. Neuropsychopharmacology : official publication of the American College of Neuropsychopharmacology 35, 870–880 (2010).

16. M. Filali, R. Lalonde, S. Rivest, Cognitive and non-cognitive behaviors in an APPswe/PS1 bigenic model of Alzheimer’s disease. Genes, brain, and behavior 8, 143–148 (2009).

17. T. Ma et al., Suppression of eIF2 alpha kinases alleviates Alzheimer’s disease-related plasticity and memory deficits. Nature Neuroscience 16, 1299–U1185 (2013).

18. S. Viana da Silva, et al., Early synaptic deficits in the APP/PS1 mouse model of Alzheimer’s disease involve neuronal adenosine A2A receptors. Nat Commun 7, 11915 (2016).

19. K. D. Harris, H. Hirase, X. Leinekugel, D. A. Henze, G. Buzsaki, Temporal interaction between single spikes and complex spike bursts in hippocampal pyramidal cells. Neuron 32, 141–149 (2001).

20. R. U. Muller, J. L. Kubie, J. B. Ranck, Jr., Spatial firing patterns of hippocampal complex-spike cells in a fixed environment. The Journal of neuroscience : the official journal of the Society for Neuroscience 7, 1935–1950 (1987).

21. J. O’Keefe, J. Dostrovsky, The hippocampus as a spatial map. Preliminary evidence from unit activity in the freely-moving rat. Brain Res 34, 171–175 (1971).

22. J. Cho, R. Bhatt, Y. Elgersma, A. J. Silva, alpha-Calcium calmodulin kinase II modulates the temporal structure of hippocampal bursting patterns. PloS one 7, e31649 (2012).

23. J. E. Lisman, Bursts as a unit of neural information: making unreliable synapses reliable. Trends Neurosci 20, 38–43 (1997).

24. M. Park et al., Chronic Stress Alters Spatial Representation and Bursting Patterns of Place Cells in Behaving Mice. Scientific reports 5, 16235 (2015).

25. L. T. Thompson, P. J. Best, Long-term stability of the place-field activity of single units recorded from the dorsal hippocampus of freely behaving rats. Brain Res 509, 299–308 (1990).

26. S. Leutgeb et al., Independent codes for spatial and episodic memory in hippocampal neuronal ensembles. Science 309, 619–623 (2005).

27. E. Bostock, R. U. Muller, J. L. Kubie, Experience-dependent modifications of hippocampal place cell firing. Hippocampus 1, 193–205 (1991).

28. R. U. Muller, J. L. Kubie, The effects of changes in the environment on the spatial firing of hippocampal complex-spike cells. The Journal of neuroscience : the official journal of the Society for Neuroscience 7, 1951–1968 (1987).

29. M. A. Wilson, B. L. McNaughton, Dynamics of the hippocampal ensemble code for space. Science 261, 1055–1058 (1993).

30. M. Padurariu, A. Ciobica, I. Mavroudis, D. Fotiou, S. Baloyannis, Hippocampal neuronal loss in the CA1 and CA3 areas of Alzheimer’s disease patients. Psychiatr Danub 24, 152–158 (2012).

31. M. A. Yassa, C. E. Stark, Pattern separation in the hippocampus. Trends Neurosci 34, 515–525 (2011).

32. S. Cayzac et al., Altered hippocampal information coding and network synchrony in APP-PS1 mice. Neurobiology of aging 36, 3200–3213 (2015).

33. C. R. Galloway et al., Hippocampal place cell dysfunction and the effects of muscarinic M(1) receptor agonism in a rat model of Alzheimer’s disease. Hippocampus 28, 568–585 (2018).

34. H. Jun et al., Disrupted Place Cell Remapping and Impaired Grid Cells in a Knockin Model of Alzheimer’s Disease. Neuron 107, 1095–1112 e1096 (2020).

35. A. J. Mably, B. J. Gereke, D. T. Jones, L. L. Colgin, Impairments in spatial representations and rhythmic coordination of place cells in the 3xTg mouse model of Alzheimer’s disease. Hippocampus 27, 378–392 (2017).

36. O. Rechnitz, I. Slutsky, G. Morris, D. Derdikman, Hippocampal sub-networks exhibit distinct spatial representation deficits in Alzheimer’s disease model mice. Curr Biol 31, 3292–3302 e3296 (2021).

37. D. Jung, Y. Huh, J. Cho, The Ventral Midline Thalamus Mediates Hippocampal Spatial Information Processes upon Spatial Cue Changes. The Journal of neuroscience : the official journal of the Society for Neuroscience 39, 2276–2290 (2019).

38. W. Skaggs, B. Mcnaughton, K. Gothard, An information-theoretic approach to deciphering the hippocampal code. Advances in neural information processing systems 5 (1992).

39. T. Indersmitten et al., In vivo Calcium Imaging Reveals That Cortisol Treatment Reduces the Number of Place Cells in Thy1-GCaMP6f Transgenic Mice. Front Neurosci 13, 176 (2019).

40. R. C. Team (2008) R: A language and environment for statistical computing. (R Foundation for Statistical Computing, Vienna, Austria).

41. M. L. Waskom, Seaborn: statistical data visualization. Journal of Open Source Software 6, 3021 (2021).

42. H. Bokil, P. Andrews, J. E. Kulkarni, S. Mehta, P. P. Mitra, Chronux: a platform for analyzing neural signals. Journal of neuroscience methods 192, 146–151 (2010).

43. M. Aguilera, V. Douchamps, D. Battaglia, R. Goutagny, How Many Gammas? Redefining Hippocampal Theta-Gamma Dynamic During Spatial Learning. Front Behav Neurosci 16, 811278 (2022).

44. L. L. Colgin et al., Frequency of gamma oscillations routes flow of information in the hippocampus. Nature 462, 353–357 (2009).

45. G. Buzsaki, Theta rhythm of navigation: Link between path integration and landmark navigation, episodic and semantic memory. Hippocampus 15, 827–840 (2005).

46. J. Okeefe, M. L. Recce, Phase Relationship between Hippocampal Place Units and the Eeg Theta-Rhythm. Hippocampus 3, 317–330 (1993).

47. M. van den Berg, D. Toen, M. Verhoye, G. A. Keliris, Alterations in theta-gamma coupling and sharp wave-ripple, signs of prodromal hippocampal network impairment in the TgF344-AD rat model. Front Aging Neurosci 15, 1081058 (2023).

48. R. A. Wirt et al., Altered theta rhythm and hippocampal-cortical interactions underlie working memory deficits in a hyperglycemia risk factor model of Alzheimer’s disease. Commun Biol 4, 1036 (2021).

49. R. Lalonde, M. Le Pecheur, C. Strazielle, J. London, Exploratory activity and motor coordination in wild-type SOD1/SOD1 transgenic mice. Brain research bulletin 66, 155–162 (2005).

50. F. L. Assini, M. Duzzioni, R. N. Takahashi, Object location memory in mice: Pharmacological validation and further evidence of hippocampal CA1 participation. Behav Brain Res 204, 206–211 (2009).

51. G. R. I. Barker, E. C. Warburton, When Is the Hippocampus Involved in Recognition Memory? Journal of Neuroscience 31, 10721–10731 (2011).

52. D. G. Mumby, S. Gaskin, M. J. Glenn, T. E. Schramek, H. Lehmann, Hippocampal damage and exploratory preferences in rats: memory for objects, places, and contexts. Learning & memory 9, 49–57 (2002).

53. A. A. Fenton et al., Unmasking the CA1 ensemble place code by exposures to small and large environments: more place cells and multiple, irregularly arranged, and expanded place fields in the larger space. The Journal of neuroscience : the official journal of the Society for Neuroscience 28, 11250–11262 (2008).

54. D. Furcila, M. Dominguez-Alvaro, J. DeFelipe, L. Alonso-Nanclares, Subregional Density of Neurons, Neurofibrillary Tangles and Amyloid Plaques in the Hippocampus of Patients With Alzheimer’s Disease. Front Neuroanat 13, 99 (2019).

55. R. de Flores, R. La Joie, G. Chetelat, Structural imaging of hippocampal subfields in healthy aging and Alzheimer’s disease. Neuroscience 309, 29–50 (2015).

56. R. A. Mease, T. Kuner, A. L. Fairhall, A. Groh, Multiplexed Spike Coding and Adaptation in the Thalamus. Cell Rep 19, 1130–1140 (2017).

57. H. L. Su, G. Alroy, E. D. Kirson, Y. Yaari, Extracellular calcium modulates persistent sodium current-dependent burst-firing in hippocampal pyramidal neurons. Journal of Neuroscience 21, 4173–4182 (2001).

58. S. Royer et al., Control of timing, rate and bursts of hippocampal place cells by dendritic and somatic inhibition. Nat Neurosci 15, 769–775 (2012).

59. C. Grienberger, X. W. Chen, A. Konnerth, NMDA Receptor-Dependent Multidendrite Ca2+ Spikes Required for Hippocampal Burst Firing In Vivo. Neuron 81, 1274–1281 (2014).

60. D. L. Hunt, N. Puente, P. Grandes, P. E. Castillo, Bidirectional NMDA receptor plasticity controls CA3 output and heterosynaptic metaplasticity. Nat Neurosci 16, 1049–1059 (2013).

61. R. Williamson, H. V. Wheal, The contribution of AMPA and NMDA receptors to graded bursting activity in the hippocampal CA1 region in an acute in vitro model of epilepsy. Epilepsy research 12, 179–188 (1992).

62. I. Bezprozvanny, M. P. Mattson, Neuronal calcium mishandling and the pathogenesis of Alzheimer’s disease. Trends Neurosci 31, 454–463 (2008).

63. A. Demuro, I. Parker, G. E. Stutzmann, Calcium signaling and amyloid toxicity in Alzheimer disease. The Journal of biological chemistry 285, 12463–12468 (2010).

64. O. Thibault, T. Pancani, P. W. Landfield, C. M. Norris, Reduction in neuronal L-type calcium channel activity in a double knock-in mouse model of Alzheimer’s disease. Biochimica et biophysica acta 1822, 546–549 (2012).

65. A. MacManus et al., Enhancement of (45)Ca(2+) influx and voltage-dependent Ca(2+) channel activity by beta-amyloid-(1-40) in rat cortical synaptosomes and cultured cortical neurons. Modulation by the proinflammatory cytokine interleukin-1beta. The Journal of biological chemistry 275, 4713–4718 (2000).

66. M. P. Mattson, Pathways towards and away from Alzheimer’s disease. Nature 430, 631–639 (2004).

67. M. P. Mattson et al., beta-Amyloid peptides destabilize calcium homeostasis and render human cortical neurons vulnerable to excitotoxicity. The Journal of neuroscience : the official journal of the Society for Neuroscience 12, 376–389 (1992).

68. C. Rovira, N. Arbez, J. Mariani, Abeta(25-35) and Abeta(1-40) act on different calcium channels in CA1 hippocampal neurons. Biochemical and biophysical research communications 296, 1317–1321 (2002).

69. E. H. Chang et al., AMPA receptor downscaling at the onset of Alzheimer’s disease pathology in double knockin mice. Proceedings of the National Academy of Sciences of the United States of America 103, 3410–3415 (2006).

70. M. Mezler, S. Barghorn, H. Schoemaker, G. Gross, V. Nimmrich, A beta-amyloid oligomer directly modulates P/Q-type calcium currents in Xenopus oocytes. Brit J Pharmacol 165, 1572–1583 (2012).

71. V. Nimmrich et al., Amyloid beta oligomers (A beta(1-42) globulomer) suppress spontaneous synaptic activity by inhibition of P/Q-type calcium currents. The Journal of neuroscience : the official journal of the Society for Neuroscience 28, 788–797 (2008).

72. N. Z. Gerges, A. M. Aleisa, L. A. Schwarz, K. A. Alkadhi, Reduced basal CaMKII levels in hippocampal CA1 region: Possible cause of stress-induced impairment of LTP in chronically stressed rats. Hippocampus 14, 402–410 (2004).

73. A. J. Silva, C. F. Stevens, S. Tonegawa, Y. Wang, Deficient hippocampal long-term potentiation in alpha-calcium-calmodulin kinase II mutant mice. Science 257, 201–206 (1992).

74. D. Y. Min et al., The alterations of Ca2+/calmodulin/CaMKII/Ca(v)1.2 signaling in experimental models of Alzheimer’s disease and vascular dementia. Neuroscience letters 538, 60–65 (2013).

75. C. A. Mackenzie-Gray Scott et al., Resilient Hippocampal Gamma Rhythmogenesis and Parvalbumin-Expressing Interneuron Function Before and After Plaque Burden in 5xFAD Alzheimer’s Disease Model. Front Synaptic Neurosci 14, 857608 (2022).

76. E. I. Moser, M. B. Moser, One-shot memory in hippocampal CA3 networks. Neuron 38, 147–148 (2003).

77. B. A. Ally, E. P. Hussey, P. C. Ko, R. J. Molitor, Pattern separation and pattern completion in Alzheimer’s disease: evidence of rapid forgetting in amnestic mild cognitive impairment. Hippocampus 23, 1246–1258 (2013).

